# Cell cortex microtubules contribute to division plane positioning during telophase in maize

**DOI:** 10.1101/2021.01.11.426230

**Authors:** Marschal A. Bellinger, Aimee N. Uyehara, Pablo Martinez, Michael C. McCarthy, Carolyn G. Rasmussen

**Affiliations:** Department of Botany and Plant Sciences, Center for Plant Cell Biology, Institute for Integrative Genome Biology, University of California, Riverside, USA; Biochemistry Graduate Group, University of California, Riverside, USA; Sonoma Technology, Inc., Petaluma, CA, USA

## Abstract

The phragmoplast is a plant-specific microtubule and microfilament structure that forms during telophase to direct new cell wall formation. The phragmoplast expands towards a specific location at the cell cortex called the division site. How the phragmoplast accurately reaches the division site is currently unknown. We show that a previously uncharacterized microtubule arrays accumulated at the cell cortex. These microtubules were organized by transient interactions with division-site localized proteins and were then incorporated into the phragmoplast to guide it towards the division site. A phragmoplast-guidance defective mutant, *tangled1*, had aberrant cortical-telophase microtubule accumulation that correlated with phragmoplast positioning defects. Division-site localized proteins may promote proper division plane positioning by organizing the cortical-telophase microtubule array to guide the phragmoplast to the division site during plant cell division.

**One Sentence Summary:** Microtubules accumulate at the cell cortex and interact with the plant division machinery to direct its movement towards the division site.

## Main Text

Cell division in plants occurs by transport of vesicles along an antiparallel microtubule array called the phragmoplast to guide new cell wall assembly (*1*). Microtubule nucleation on pre-existing microtubules promotes phragmoplast growth towards the cell cortex (*2*–*4*). How the phragmoplast is directed towards a specific cortical location, called the division site, is still unknown (*5, 6*). In land plants, the future division site location can be accurately predicted by a microtubule structure that assembles in G2 at the cell cortex called the preprophase band (PPB). Several proteins co-localize with the PPB, and then remain at the division site until division is completed. These division-site-localized proteins are thought to promote phragmoplast guidance to the division site because phragmoplasts in corresponding mutant cells often do not return to the division site (*7*–*10*). Several division-site-localized proteins are microtubule- or microfilament-bundling or motor proteins (*7, 11*–*13*) suggesting that division site positioning may be mediated by local alterations in cytoskeletal dynamics. Current models propose that division-site and phragmoplast-localized proteins pull or push cytoskeletal filaments within the phragmoplast to guide it to the division site. These models propose that microtubules attached to and nucleated from the phragmoplast, “peripheral microtubules”, interact with division site localized proteins and actin filaments to guide the phragmoplast (*11, 12*). Microtubules and microtubule nucleators such as gamma tubulin have long been observed at the cell cortex during telophase in monocots and dicots but their function is unknown e.g. (*14*–*17*). A previously proposed function of cortical-telophase microtubules is to prepopulate the cortex for microtubule reorganization during G1 (*18*), but here we demonstrate that cortical telophase microtubules are organized by transient interaction with the division site-localized protein TAN1 and direct the movement of the phragmoplast towards the division site.

Live-cell imaging of symmetrically dividing maize leaf epidermal cells during telophase revealed an unexpected population of cell-cortex-localized microtubules that were independent from the phragmoplast (Fig. S1). Cortical-telophase microtubules were nucleated from the cell cortex during the anaphase to telophase transition and were present in over 90% of wild-type cells during telophase (n =173/190 cells from 26 plants, Fig. 1A). Cortical-telophase microtubule arrays covered 33 +/-2% (mean +/-SEM) of the cell cortex (Fig. 1C) with an average anisotropy of 0.12 +/-0.01 arbitrary units (Fig. 1B). Anisotropy values reflecting relative orientation of cortical-telophase arrays were similar to interphase shoot-meristem Arabidopsis microtubule arrays (*19*). The cortical-telophase microtubules were typically arranged into anti-parallel arrays perpendicular to the division site (∼50% within 10 degrees of perpendicular, n = 38 microtubule arrays from 19 cells, 7 plants, Fig.1D) with their plus-ends facing the division site. We also observed telophase cortical microtubule arrays in Arabidopsis root cells (Fig. S2) similar to previous reports on microtubule-nucleating protein accumulation at the cell cortex e.g. (*14, 20*) or cortical microtubules in moss e.g. (*11*). Therefore, cortical-telophase microtubule arrays were abundant and oriented perpendicular to the division site in maize epidermal cells and may be a conserved feature of plant cells.

**Fig 1.**
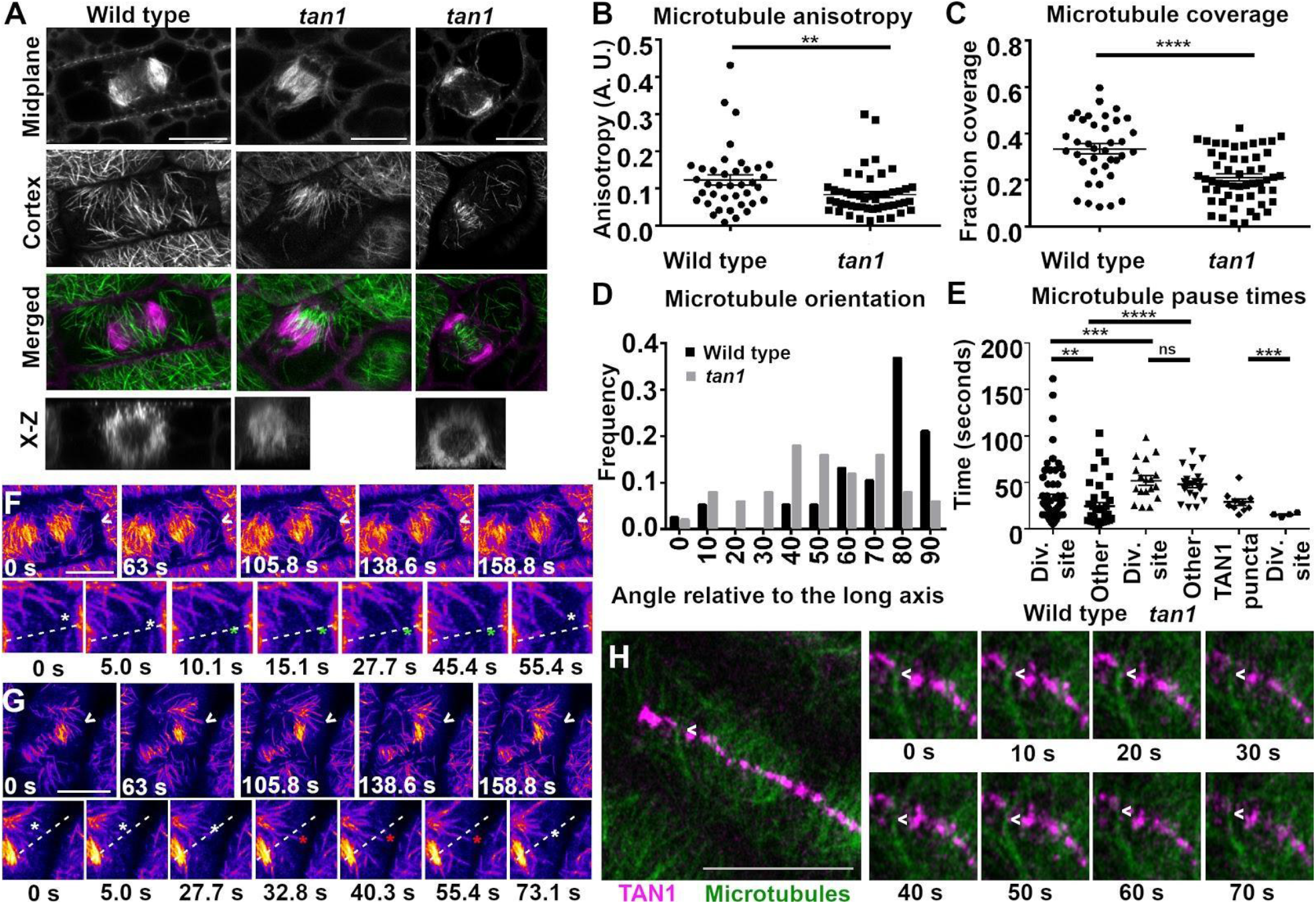
Cortical-telophase microtubules were oriented by interaction with division-site localized proteins. A) A wild-type cell with cortical-telophase microtubules (left), *tangled1* (*tan1)* mutant cell with uneven (middle) or sparse cortical-telophase microtubules (right). Cortical-telophase microtubule array (B) anisotropy and (C) relative coverage was significantly higher in wild-type (38 arrays) than *tan1* (50 arrays) cells. D) Histogram of cortical-telophase microtubule array angles. E) Dot plot of microtubule plus-end pause times at indicated locations in wild-type (left) and *tan1* mutants (middle) and including pause at TAN1 puncta and other regions of the division site without TAN1 puncta in wild-type cells. F and G) phragmoplast (orange, false-colored) and cortical telophase microtubules (pink-purple) interaction with the division site (arrowhead (top) and dashed line (bottom)). F) Wild-type microtubule (white asterisk) pausing (green asterisk) at the division site (dashed line) in the bottom panel (scaled 250%). G) *tangled1* telophase cell bottom panel (scaled 148%) white asterisk indicates a cortical microtubule growing and shrinking. The red asterisk shows a microtubule passing through the division site. H) Microtubule (CFP-TUBULIN, green) pausing near TAN1-YFP puncta (magenta) from 10 s to 20 s, then shrinking from 30 s to 40 s, growing then pausing again at 60 s. Bar = 10 µm. Significance measured with Mann-Whitney U test. P-values ns not significant, ** <0.01, *** <0.001, **** <0.0001.

Aberrant cortical-telophase microtubule arrays were observed in a mutant with defects in division plane orientation, *tangled1 (tan1)*. TAN1, a microtubule-binding protein that localizes to the division site, is required for proper phragmoplast guidance to the division site (*8, 21, 22*). TAN1 crosslinks and bundles microtubules in vitro and co-localizes with microtubules in vivo (*13*). We hypothesized that loss of TAN1 from the division site would lead to defects in cortical-telophase microtubule organization. Cortical-telophase microtubule arrays were sparse or missing in nearly 30% of *tan1* mutant cells (n = 24/122 cells from 24 plants, e.g. Fig. S3). When cortical-telophase arrays were present in *tan1* mutant cells, they were often unevenly distributed or less dense (Fig. 1A and 1C). Further, cortical-telophase microtubule arrays were both less anisotropic (Fig.1B) and not typically oriented toward the division site (Figure 1D, median orientation 49.5 +/-3 degrees, P < 0.0001 Mann-Whitney test) compared to wild-type arrays. These data suggest that TAN1 promotes proper cortical-telophase microtubule array organization.

To understand how telophase cortical microtubules formed perpendicular arrays with their plus-ends facing the division site, we examined individual microtubules interacting with the division site. In wild-type cells, microtubule plus-ends were transiently stabilized by pausing at the division site (Fig. 1E, 1F, Movie S1). When cortical-telophase microtubules contacted the division site, 77% of microtubules paused (n = 69/90 microtubules, 5 cells, 3 plants, Table S1), 13% underwent immediate catastrophe after touching the division site, and 9% passed through the division site without alteration of their trajectories. Median pausing time was 33.4 seconds at the division site but 13.0 seconds in other locations. Transient stabilization of microtubule plus-ends at the division site may promote overall perpendicular orientation. In contrast to wild-type cells, cortical-telophase microtubules were not transiently stabilized at the division site in *tan1* mutants, showing no significant difference in microtubule pausing at the division site versus in other cortical locations (Fig. 1E), although generally microtubule pause times were increased (median 49.1 +/-5.0 seconds, n = 38). The majority of cortical-telophase microtubules in the *tan1* mutant grew past the division site without any alteration in their trajectories (76% n = 29/38, example in Fig.1G, Table S1), instead of pausing (24% n = 9/38) or shrinking (0%, n = 0/38). This suggests that division site localized TAN1 directly or indirectly promotes both microtubule plus-end pausing and shrinking at the division site.

Cortical-telophase microtubules paused near TAN1-YFP puncta longer than other regions of the division site. TAN1 forms discrete puncta at the division site during telophase (Fig.1H) (*8, 21, 23*), and transiently crosslinks microtubules in vitro (*13*). We measured microtubule pause time at the division site. Microtubule plus-ends paused near TAN-YFP1 puncta for ∼26 seconds (n = 10 microtubules). In contrast, microtubule plus-ends that contacted regions of the division site without TAN1-YFP puncta, paused for ∼15 seconds (n = 4 microtubules, Fig. 1E). Together, this suggests that TAN1 or other division-site localized proteins in close proximity, may promote cortical telophase microtubule plus-end pausing. This microtubule interaction is consistent with in vitro dynamic assays where TAN1 transiently crosslinked microtubules (*13*).

Cortical-telophase microtubules interacted dynamically with the division site, leading to perpendicular arrays, but they were also added into the phragmoplast as the phragmoplast expanded towards and along the division site. We assessed how individual microtubules from the cortical-telophase array interacted with the phragmoplast by time-lapse imaging. When cortical-telophase microtubules contacted the phragmoplast, most (78%, n = 197/252 microtubules from 5 cells from 3 plants) were incorporated into the phragmoplast by parallel bunding (Fig. 2A, Movie S2). After bundling into the phragmoplast, the microtubules would either remain connected to the original cortical-telophase array during the 252 second timelapse (41%, n = 103/252), or become fully incorporated into the phragmoplast by severing the connection between the cortical-telophase array and the phragmoplast (37%, n = 94/251, Fig. 2B, Movie S3). Microtubule severing sometimes occurred at the distal end of the phragmoplast (for a model see Fig. S4). We speculate that severing was performed by the microtubule severing protein KATANIN localized to the distal phragmoplast via the microtubule-binding protein, MACET4/CORD4 (*24, 25*). The remaining telophase cortical microtubules that contacted the phragmoplast underwent catastrophe after touching the phragmoplast (22%, n = 55/252, Fig. 2C, Movie S4). Most (66%, n = 166/252) telophase cortical microtubules interacted with the leading edge, although others interacted with the lagging edge (n = 86/252) and then primarily were incorporated into the phragmoplast by low-angle parallel bunding (Table 1).

**Fig 2.**
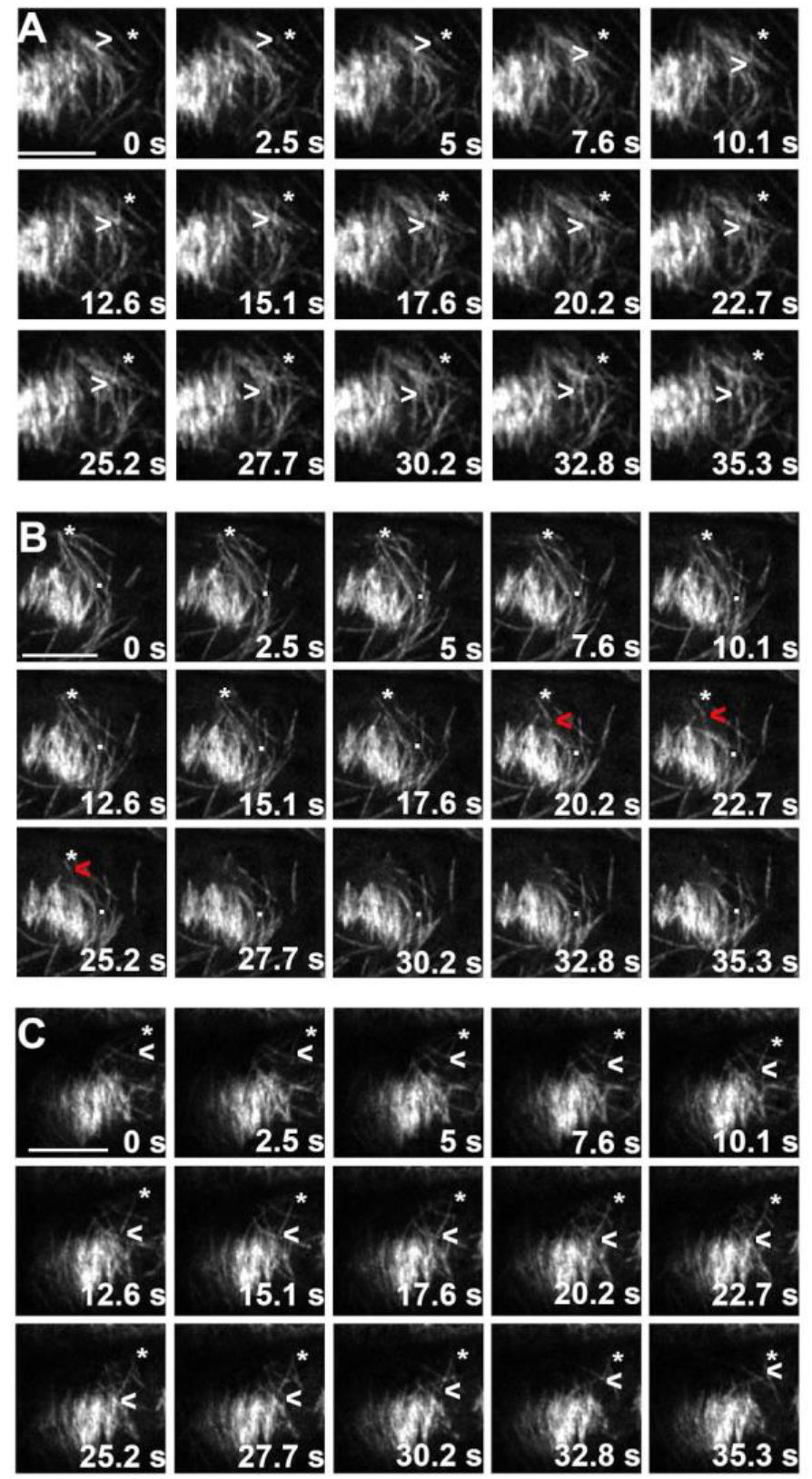
Timelapse images showing cortical-telophase microtubules interacting with the phragmoplast. Asterisks indicate the cortical-telophase microtubule minus-ends, white arrowheads show growing and shrinking plus-ends, white dots show stable cortical-telophase plus-ends, and red arrowheads show severing followed by depolymerization A) A microtubule was incorporated into the phragmoplast by parallel bundling. B) Severing of a microtubule from the cortical-telophase array led to incorporation of the microtubule into the phragmoplast. C) A microtubule encountered the phragmoplast, and then underwent catastrophe without incorporation into the phragmoplast. Bars are 5 µm, Time was rounded to a tenth of a second.

**Table 1.**
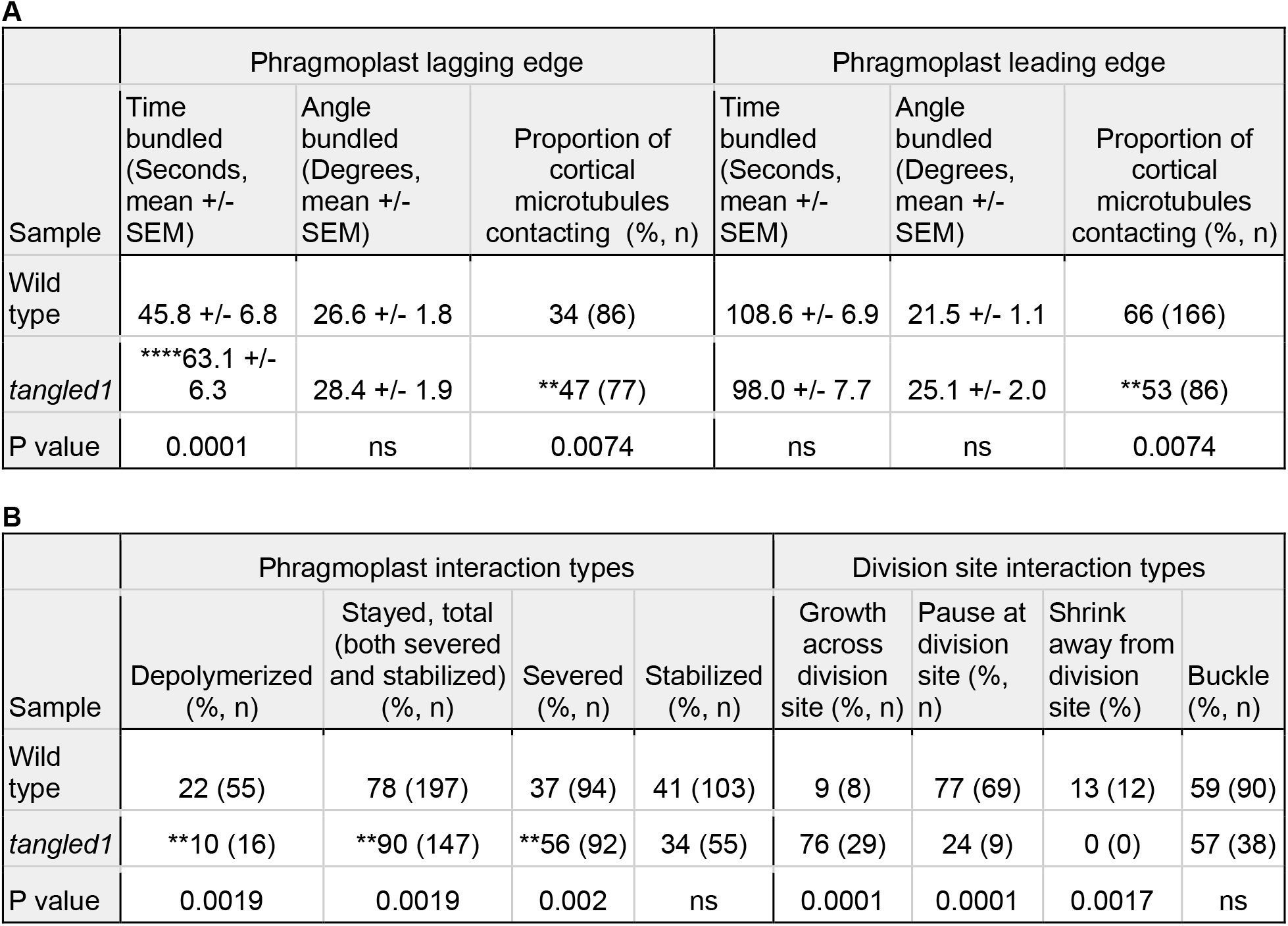
Quantification of individual interaction and bundling events between cell cortex telophase derived microtubules and the phragmoplast. (A) Summary of cell cortex derived microtubule bundling times and angles with the phragmoplast. (B) Summary of cell cortex derived microtubule interact types with the phragmoplast or putative division site. (B) Fisher’s exact-test was used and significant differences were indicated by (**) P < 0.01, (****) P < 0.0001. Phragmoplast interacting MTs: WT (n = 252, 5 cells, 3 individuals), *tan1* (n = 163, 5 cells, 3 individuals). Division site interacting MTs: WT (n = 90), *tan1* (n = 38).

Cortical-telophase microtubules, when present, were also added into *tan1* mutant phragmoplasts. Similar to wild-type microtubules, *tan1* cortical-telophase microtubules were incorporated into the phragmoplast, although relatively more microtubules interacted with the lagging edge of the phragmoplast (Table S1). Proportionally more of the microtubules that interacted with the phragmoplast were eventually incorporated in *tangled1* mutant phragmoplasts (90% n = 147/163 versus 78% in wild-type cells n =197/252 Table 1). Therefore, cortical telophase microtubules in close contact with the phragmoplast were primarily added into the leading edge in both wild-type and *tan1* mutant cells.

Finally, cortical-telophase microtubule accumulation predicted the direction of phragmoplast movement. We hypothesized that the asymmetric accumulation of cortical-telophase microtubules on one side of the phragmoplast would lead local microtubule addition on the corresponding side of the phragmoplast followed by movement of the phragmoplast towards that direction of the cell, while more microtubules on the opposite side would promote phragmoplast movement towards the opposite direction of the cell. We compared the phragmoplast trajectory with the relative accumulation of cortical-telophase microtubules “above” and “below” the phragmoplast using timelapse imaging (Fig. 3A-C). The phragmoplast trajectory was measured as an angle parallel to the division site (Fig. 3A): positive angle values indicate that the phragmoplast angle moved above the division site. Two equally-sized region-of-interest boxes were selected (Fig. 3B) and relative cortical-telophase microtubule accumulation above and below the phragmoplast was measured by subtracting below from above microtubule coverage. Positive values indicate that more microtubules accumulate above the phragmoplast. Phragmoplast expansion direction in wild-type cells typically followed a flat trajectory within 5 minutes, with < 10 degree overall change (n= 5, Fig. 3D, Fig. S5). During longer timeframes (18-30 minutes), wild-type phragmoplast trajectories were more variable, but overall, did not persistently change direction (n = 2/4, Fig. S6), consistent with previous timelapse observations (*8*). In wild-type cells with little overall phragmoplast angle displacement, cortical-telophase microtubule accumulation varied over time, but did not maintain uneven accumulation (Fig. 3E, S6B and S6D). In contrast, sustained uneven accumulation of telophase cortical microtubules was correlated with phragmoplast movement in the same direction (Fig. 3F, Fig. S6A and S6C). In *tangled1* mutants, both phragmoplast expansion direction and cortical-telophase microtubule array accumulation were more variable (Fig S7). Over longer timeframes, uneven sustained cortical-telophase microtubule accumulation in *tan1* mutants also correlated with changes in phragmoplast trajectories in the same direction (Fig. 3G, Fig S8).

**Fig 3.**
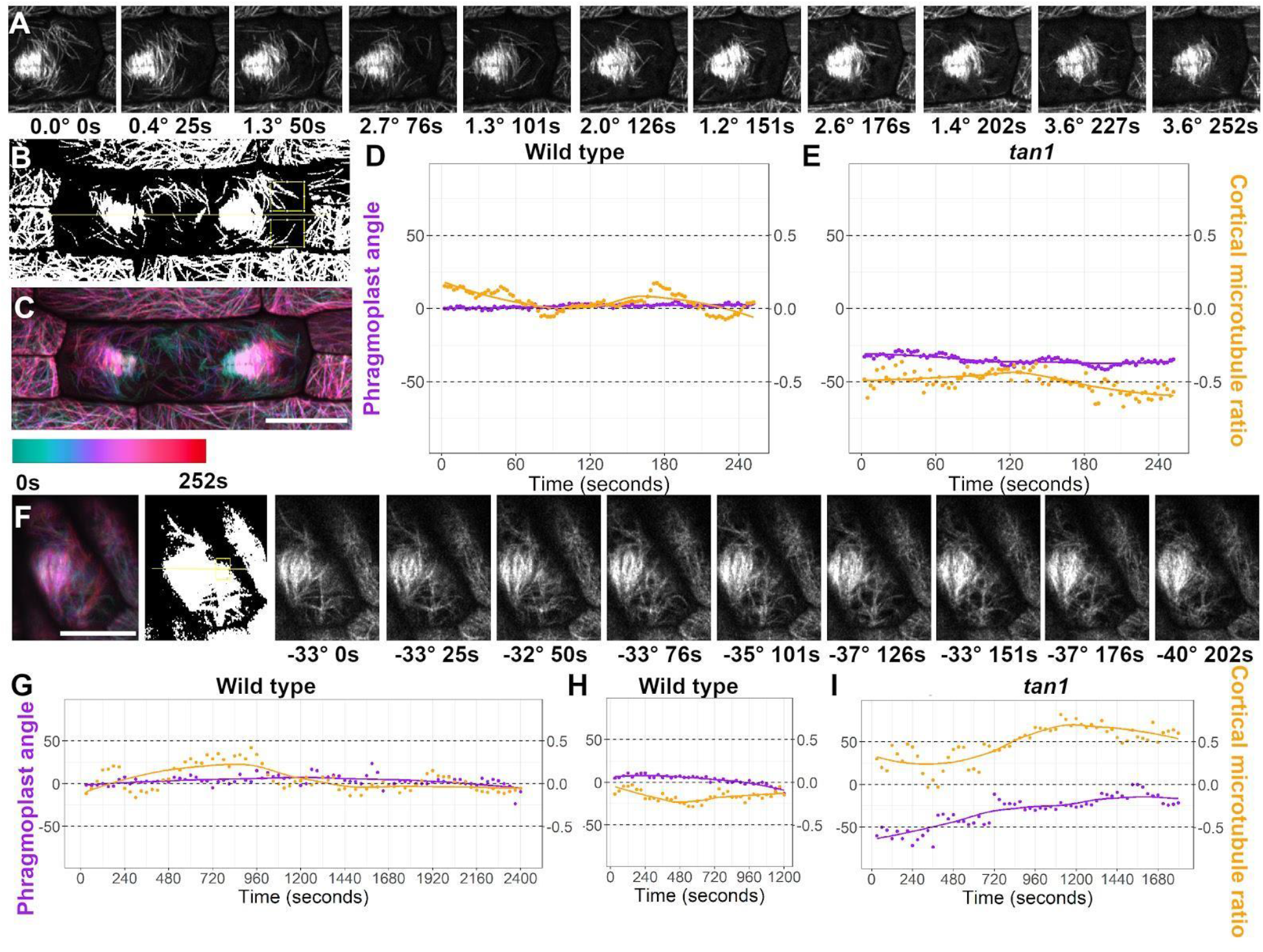
Long-term uneven accumulation of cortical-telophase microtubules was correlated with changes in phragmoplast direction. A - D) A wild-type phragmoplast A) Timelapse with seconds and phragmoplast angle parallel to the division site indicated below. B) Thresholded image with regions of interest (yellow rectangles) selected to measure relative cortical-telophase microtubule accumulation above and below the phragmoplast. The division site is indicated by a yellow line. C) Time-projection with time-color legend. D) Graph comparing phragmoplast angle changes over time (purple) and relative cortical-telophase microtubule accumulation (orange) above or below the phragmoplast. E-F) A *tan1* phragmoplast E) Graph of *tan1* phragmoplast angle and cortical-telophase microtubule changes over time. F) Timelapse of *tan1* phragmoplast angle and time below. G-I) Longer timelapses: G) A wild-type cell with little overall phragmoplast movement. H) wild-type cell with consistent cortical-telophase microtubule accumulation below the phragmoplast and phragmoplast angle movement down I) *tan1* cell with consistent cortical-telophase microtubule accumulation above the phragmoplast with phragmoplast angle movement towards the top. Bars = 10 µm.

We showed that cortical-telophase microtubule arrays accumulated perpendicular to the division site, due to transient microtubule plus-end pausing at the division site, and were often added into the phragmoplast by parallel bundling. Uneven cortical-telophase microtubule accumulation was correlated with changes in phragmoplast trajectories. These data suggest that cortical telophase microtubules are incorporated into the phragmoplast to fine-tune positioning such that the phragmoplast reaches the division site. Microtubule and microfilament interactions with division-site-localized proteins such as MyosinVIII are likely conserved plant division plane positioning features suggesting their combined action together with cortical-telophase microtubules guides the phragmoplast (*11*). Cytoskeleton-mediated division-plane corrections also occur during telophase in mouse epithelial cells suggesting that analogous mechanisms occur in other eukaryotes (*26*)

## Materials and Methods

### Plant growth and imaging conditions

Maize plants were grown in 1L pots in standard greenhouse conditions. Maize plants between 3 and 5 weeks old containing YFP-TUBULIN, CFP-TUBULIN, TANGLED1-YFP (*27, 28*) or the *tangled1* mutant were used for imaging and identified by microscopy or by genotyping as previously described (*8*). Leaves were removed until the ligule height was <2 mm. Adaxial symmetrically dividing leaf blade samples were mounted in water between cover glass and glass slides (Fisherbrand) or in a Rose chamber, as previously described (*29*). Three or more plants of each genotype were analyzed. Room temperatures during imaging were between 21 and 24 °C.

Arabidopsis plants were grown on ½ strength Murashige and Skoog (MS) media solidified with 0.8% Agar. Plates sealed with surgical tape (3M) and grown vertically in a growth chamber (Percival) at 24 C, 16-h light, 8-h dark. Arabidopsis plants between 3 and 5 days after germination were imaged. Arabidopsis plants with CFP-TUBULIN were identified by microscopy. Seedlings were mounted in water. Root epidermal cells from the meristematic zone were imaged at 23 °C.

### Confocal microscopy

Micrographs and short timelapses were taken with a Zeiss LSM 880 equipped with Airyscan with a 100X, NA = 1.46, oil immersion objective lens. A 514 excitation laser with bandpass (BP) filters 465-505 with longpass (LP) 525 filter was used with Airyscan superresolution mode. Images captured using the Zeiss LSM 880 were processed using default Airyscan settings with ZEN software (Zeiss). For longer time lapse imaging, with 30 second intervals, a Yokogawa W1 spinning disk microscope with EM-CCD camera (Hamamatsu 9100c) and Nikon Eclipse TE inverted stand was used with a 100x, NA 1.45, oil immersion objective lens controlled with Micromanager-1.4 with ASI Piezo Z stage and 3 axis DC servo motor controller. Solid-state lasers (Obis) between 40 to 100 mW were used with standard emission filters from Chroma Technology. For YFP-TUBULIN or TANGLED1-YFP, a 514 laser with emission filter 540/30 was used. For CFP-TUBULIN a 445 laser with emission filter 480/40 was used. For the membrane dye FM4-64, the 516 nm laser with emission filter 620/60 was used.

Telophase cells were identified by the presence of a phragmoplast, telophase cortical microtubules were imaged on the cortical edge of epidermal cells. Two-dimensional projections, time projections and three dimensional reconstructions of Z stacks and time-lapse images were generated in FIJI (ImageJ, http://rsb.info.nih.gov/ij/). Image brightness and contrast was altered using the linear levels option, and figures were assembled with FIJI and Gnu Image Manipulation Program (GIMP https://www.gimp.org/downloads/). Drift during timelapse was corrected with StackReg https://imagej.net/StackReg using the translation option (*30*).

### Quantification of microtubule array organization and coverage

Maize lines expressing YFP-TUBULIN were used to examine the microtubule cytoskeleton. To measure anisotropy (Figure 1B) and orientation (Figure 1D), TIFF image files were converted to PNG files using Fiji software and processed with the FibrilTool plugin (*19*). For wild-type plants, 38 arrays from 19 transverse cell divisions during telophase were measured from 5 plants with median anisotropy 0.11 +/-0.01 A.U. For *tan1* mutants, 50 arrays from 25 transverse cell divisions during telophase were measured from 9 plants (0.07 +/-0.01 A.U.). Differences in anisotropy were analyzed with Graphpad Prism and statistical significance was determined with a Mann-Whitney *U*-test, P = 0.0054.

To measure percent microtubule coverage in Figure 1C, image files were made binary and thresholded using mean fluorescence and processed using the Area/Volume fraction function in the BoneJ plugin (https://imagej.net/BoneJ, (*31*). The median value for wild type cells (n = 38 arrays, coverage fraction 0.33 +/-0.02) is significantly different from median value for *tangled1* mutant cells (n = 50 arrays, coverage fraction 0.20 +/-0.01, Mann-Whitney test, P < 0.0001.) Timelapse imaging was used to measure microtubule interactions at the division site, near TAN1 puncta, and with the phragmoplast. The division site was defined as a location either parallel or perpendicular to the long axis of the cell and on the phragmoplast midplane position. In *tan1* mutants, the “division site” was defined the same way unless the phragmoplast was misoriented, in which case the “division site” was defined only by the midplane of the phragmoplast. Individual microtubule movements were measured, binned into categories (for phragmoplast interactions) and graphed in GraphPad Prism and statistically analyzed with Mann-Whitney test.

Timelapse imaging was used to compare the abundance of cortical-telophase microtubules and the phragmoplast angle over time. The phragmoplast angle at each timepoint was measured in FIJI, and saved in Google sheets or Excel (Microsoft Office). Timelapse image files were first processed to remove drift using the transformation selection within StackReg, and to correct for photobleaching using bleach correction (exponential fit) in FIJI. Next, images were made binary and thresholded using mean fluorescence and processed using the Area/Volume fraction function in the BoneJ plugin (*31*), Two equally-sized regions-of-interest (ROIs) were selected above and below the phragmoplast, such that the ROIs captured cortical-telophase microtubule accumulation near the phragmoplast but not touching the phragmoplast at any time frame. The bottom ROI was subtracted from the top ROI in google sheets. If the value was positive, it indicated more microtubules on the top of the phragmoplast. Both phragmoplast angle and relative cortical-telophase microtubule accumulation were graphed together by time using R, RStudio (Version 1.3.1093) and ggplot2 (*32, 33*). Figures were made using Gnu Image Manipulation Program (Gimp, version 2.10.22) using no interpolation during scaling and linear adjustments to levels.

## Acknowledgments

Thanks to Professor Meng Chen (UCR) for procuring funds for the Zeiss 880 (3R01GM087388-06S2) and Dr. David Carter for Zeiss 880 training.

## Author contributions

Marschal A. Bellinger: Formal Analysis, Investigation, Writing – Original Draft, Visualization. Aimee N. Uyehara: Formal analysis, Investigation, Writing – Review and Editing. Pablo Martinez: Investigation, Writing – Review and Editing. Michael McCarthy: Resources, Visualization. Carolyn Rasmussen: Formal analysis, Investigation, Writing – Original Draft, Visualization, Supervision, Project administration, Funding acquisition.

## Data and materials availability

All data are available in the main text or the supplementary materials. Materials are at the maize cooperative (http://maizecoop.cropsci.uiuc.edu/) or are available by request.

## Supplementary Materials

Figures S1-S8

Movies S1-S4

**Fig S1:**
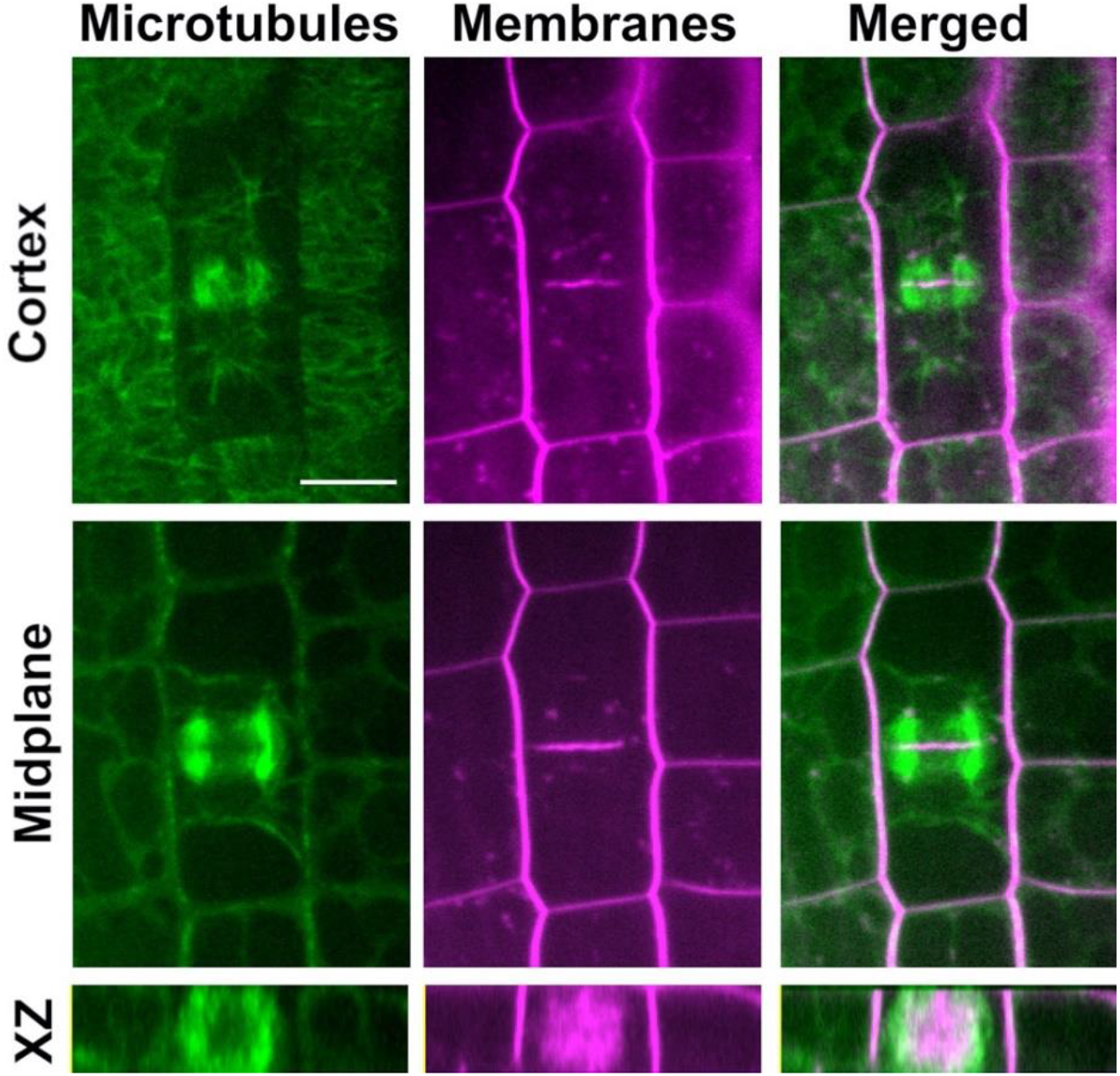
Cortical telophase microtubules were distinct from phragmoplast microtubules. Microtubules labeled with YFP-TUBULIN in green, membranes labeled with FM4-64 in magenta. Merged image shows both. Midplane indicates the middle of the cell, XZ image is a rotated projection along the XZ axis of the phragmoplast. Bar is 10 μm.

**Fig S2.**
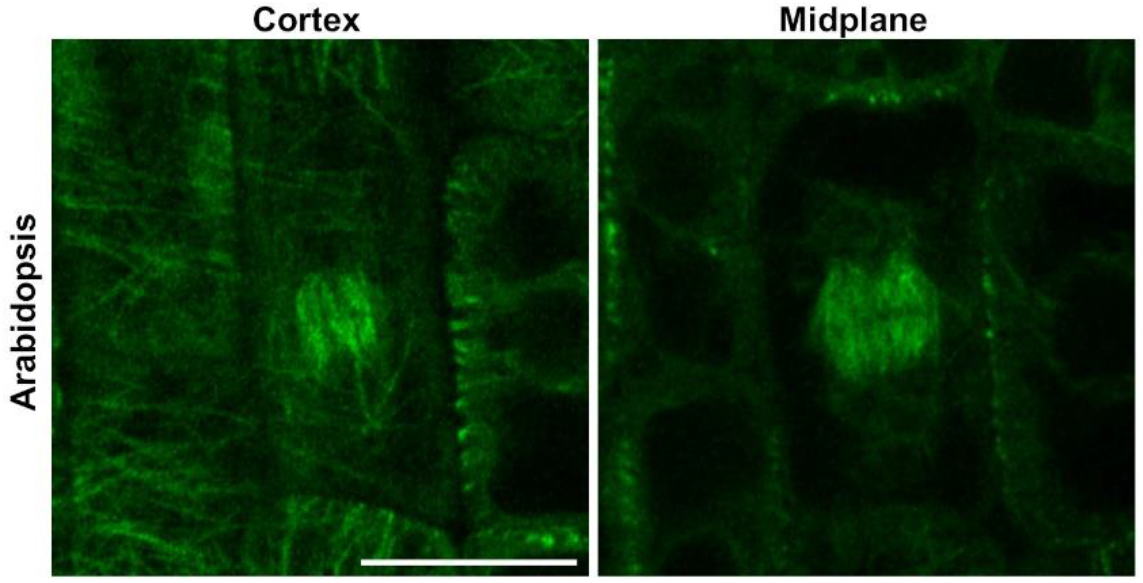
Cortical telophase microtubules in *A. thaliana* root cell with a phragmoplast. Bar is 10 μm.

**Fig S3.**
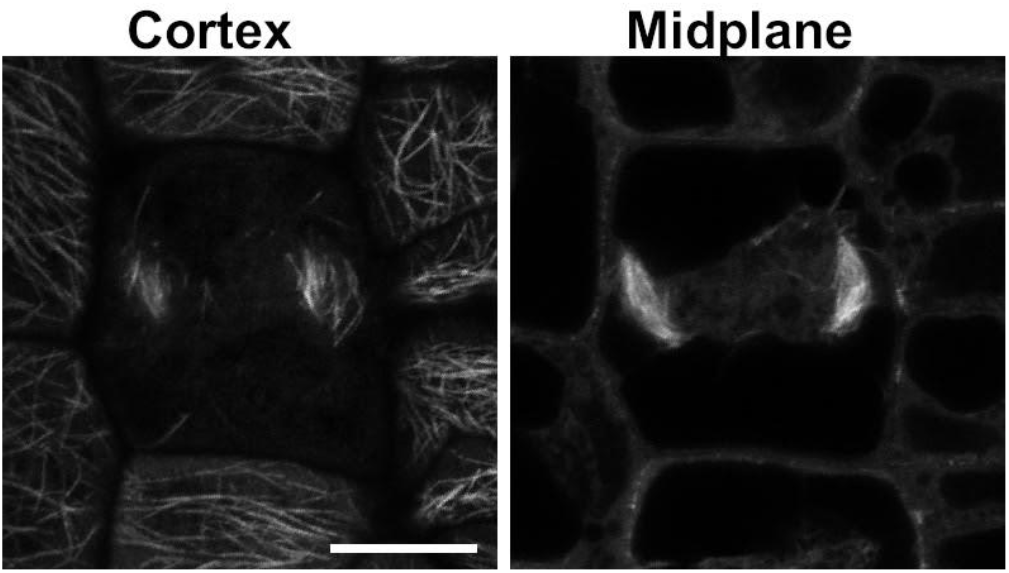
*tan1* mutant cell in telophase with a sparse cortical telophase microtubule array. Bar is 10 μm.

**Fig S4.**
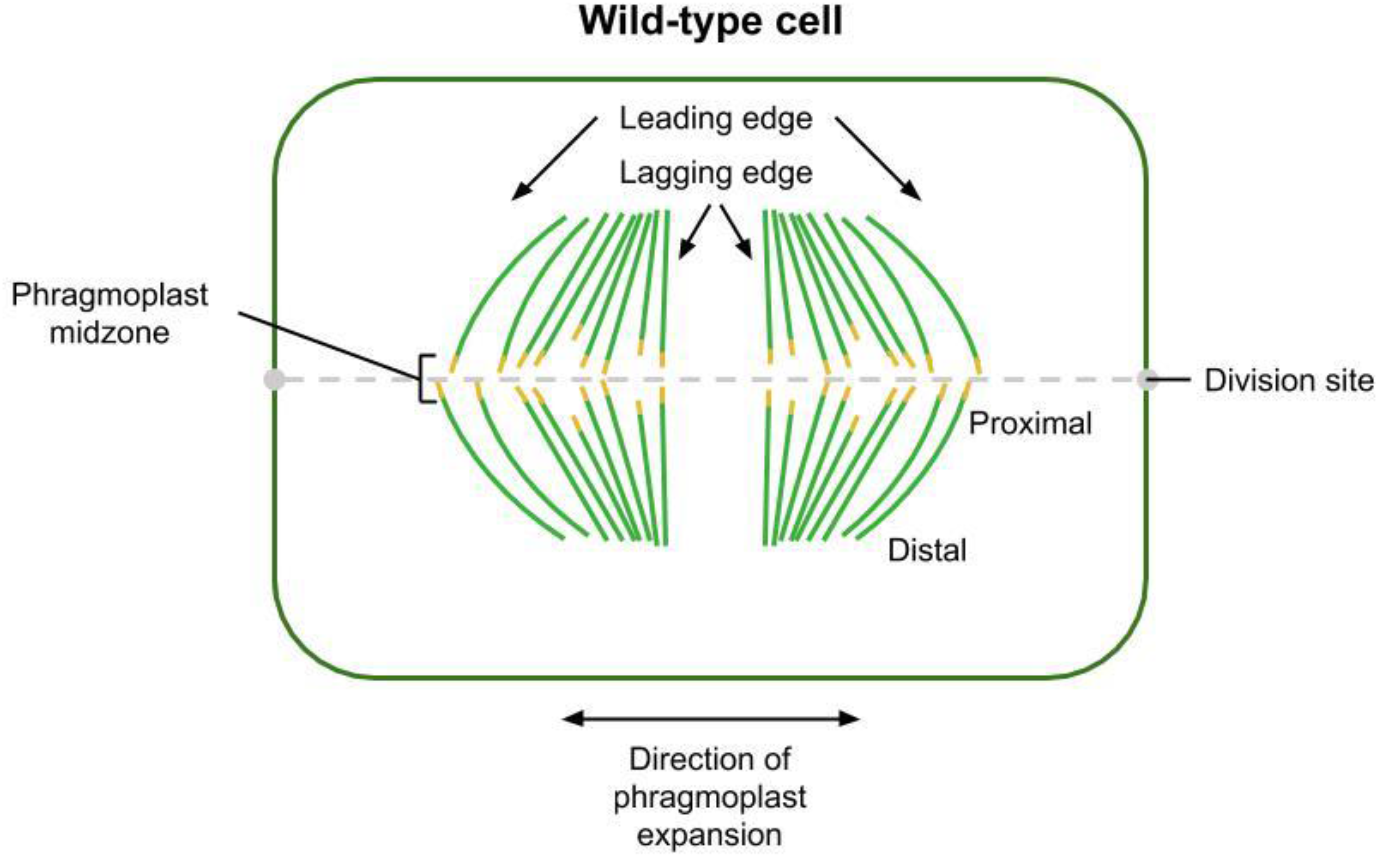
Attributes of a wild-type phragmoplast. The phragmoplast, microtubules (green), plus-ends (yellow), expands towards the division site. The outer and inner region of the phragmoplast are labeled the leading and lagging edges respectively. The plus ends are near the proximal side, while the minus ends are near the distal side.

**Fig S5.**
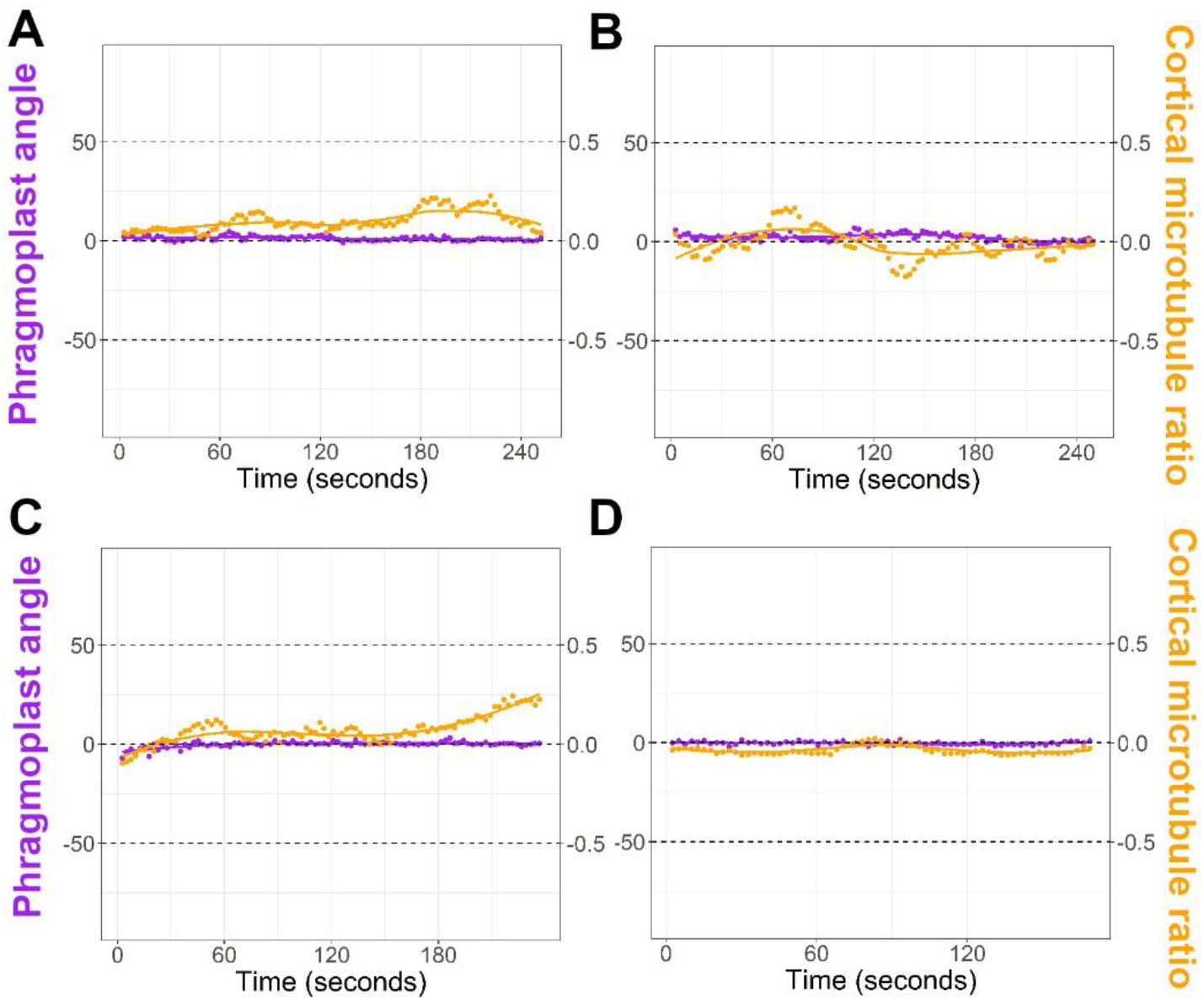
Short time-lapses (<5 minutes) of four different wild-type phragmoplasts showing little change in direction.

**Fig S6.**
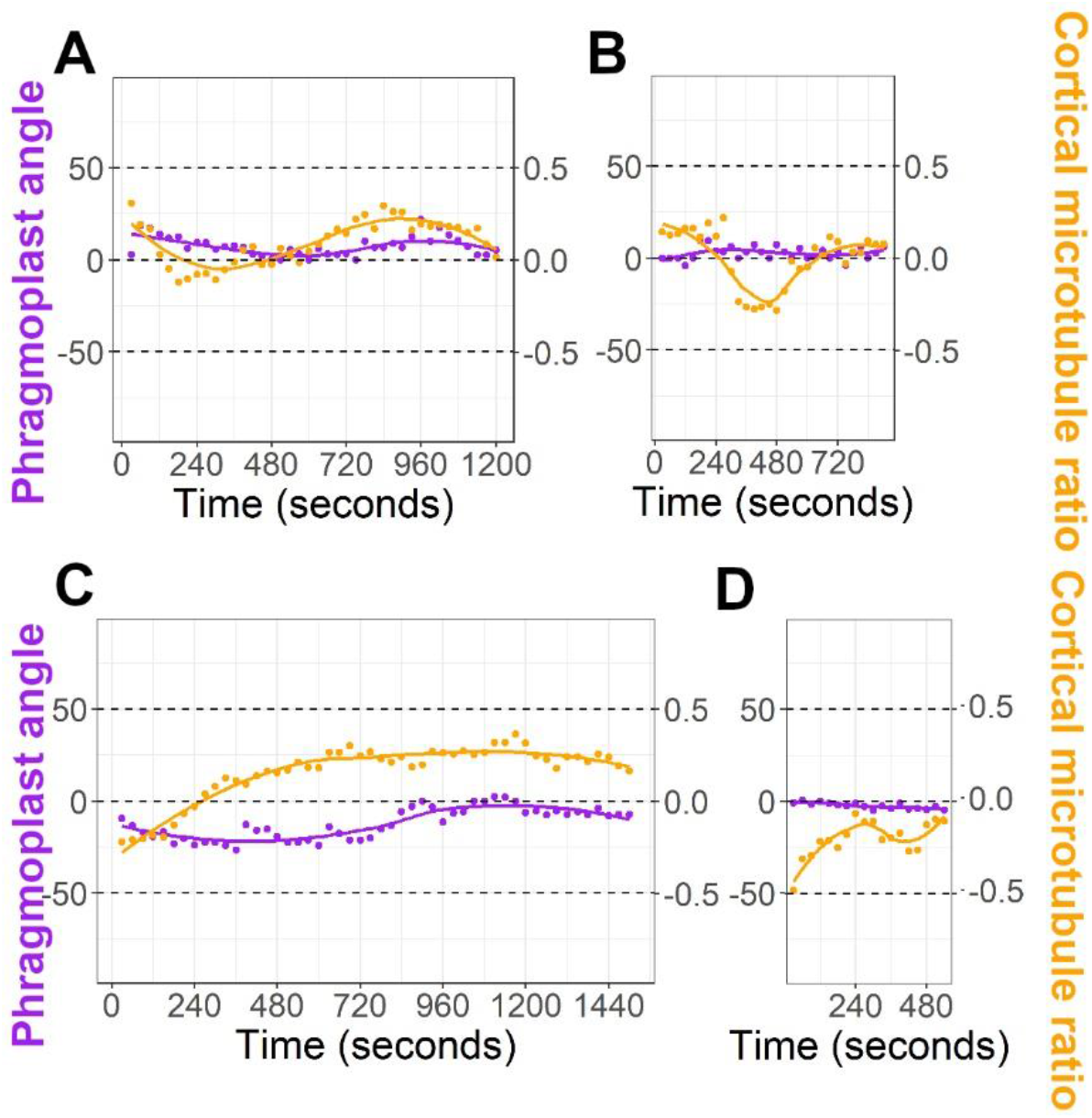
Longer timelapses (>8 minutes) of wild-type phragmoplasts with >10 degree changes in the direction of movement (A, C) and (B, D) with < 10 degree changes in the direction of movement.

**Fig S7.**
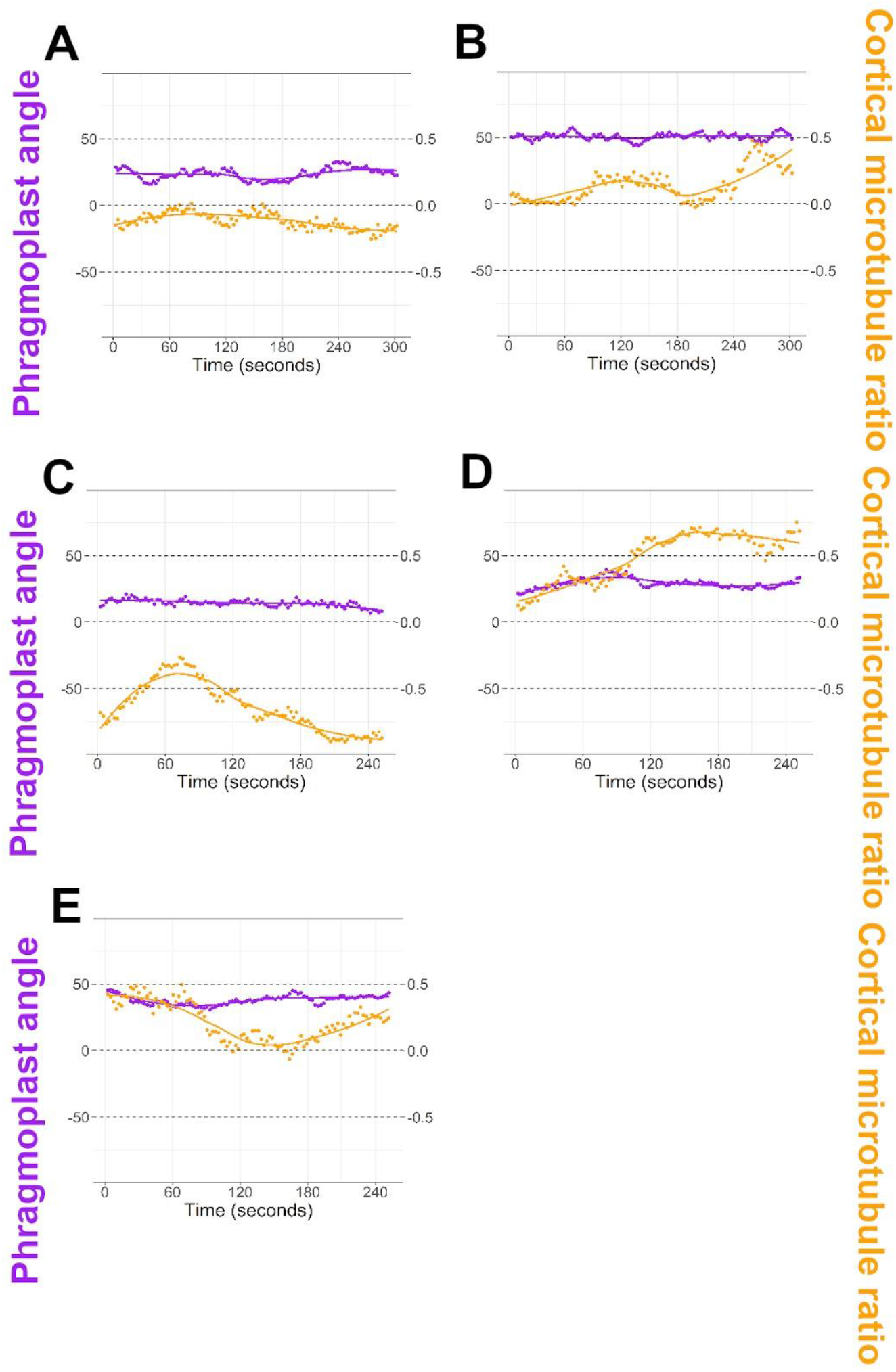
Short timelapses (5 minutes or less) of five different *tan1* phragmoplasts showing little change in direction, but frequently unevenly distributed cortical-telophase array.

**Fig S8.**
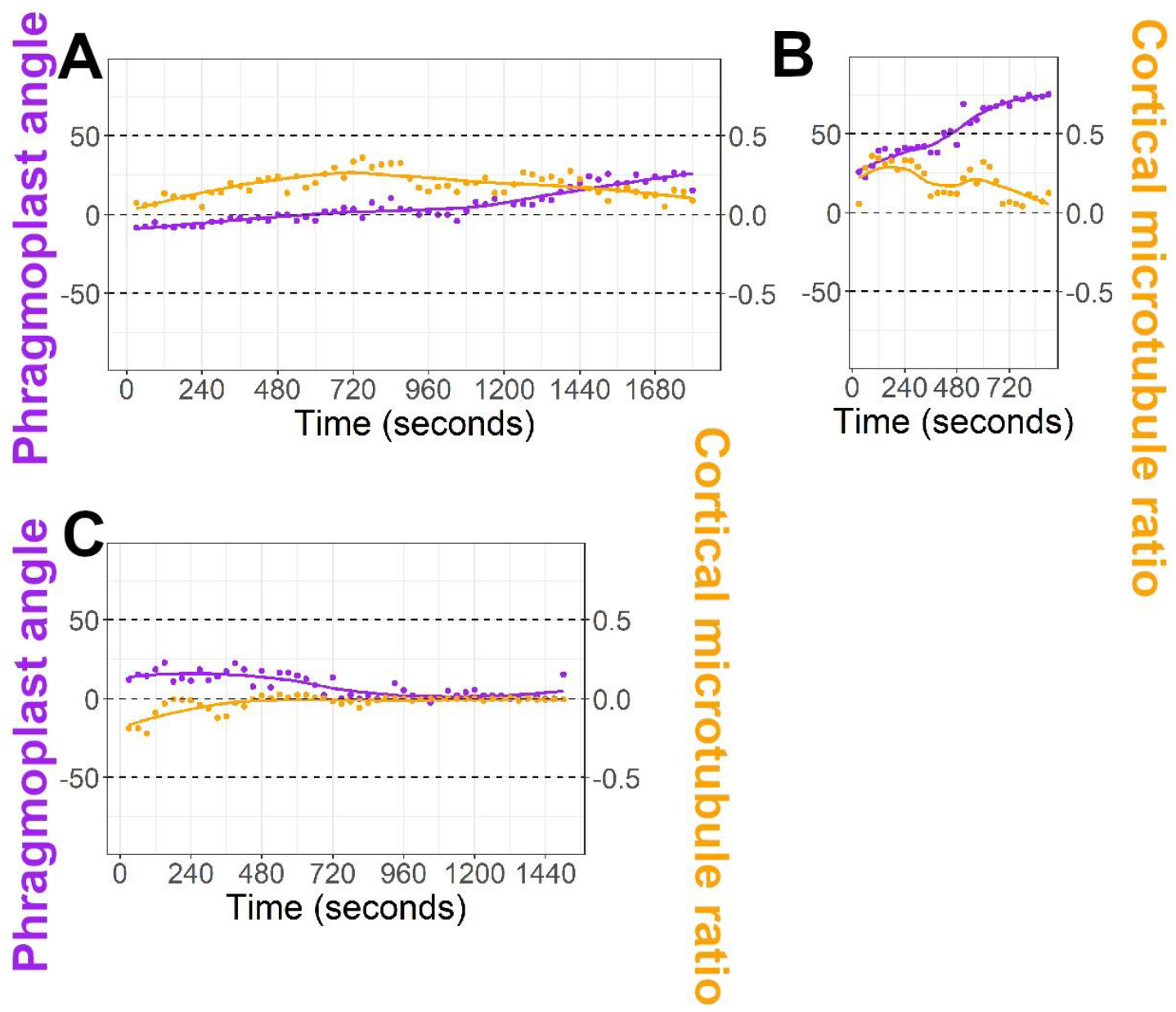
Longer timelapses (>12 minutes) of *tan1* phragmoplast angle measurements compared to cortical microtubule ratio. All show >10 degree phragmoplast angle changes.

**Movie S1**. Cell cortex of a maize epidermal cell in telophase showing cortical-telophase microtubules interacting with the phragmoplast. Time indicated is in seconds.

**Movie S2**. Cortical-telophase microtubule with minus end indicated with a white dot, plus end indicated with a yellow dot showing parallel bundling into the phragmoplast. Time indicated in seconds.

**Movie S3**. Cortical-telophase microtubule with minus-end indicated with a white dot and plus end indicated with a yellow dot. Severing location indicated with a red dot, which follows the new minus end (as it is incorporated into the phragmoplast), and a green dot, showing the new plus end of the cortical-telophase microtubule undergoing catastrophe (shrinking). Time indicated in seconds.

**Movie S4**. Cortical-telophase microtubule with minus end indicated with a white dot, and plus end indicated with a yellow dot showing the microtubule touching the phragmoplast, then undergoing catastrophe (shrinking). Time indicated in seconds.

## References

1. A. Smertenko, F. Assaad, F. Baluška, M. Bezanilla, H. Buschmann, G. Drakakaki, M.-T. Hauser, M. Janson, Y. Mineyuki, I. Moore, S. Müller, T. Murata, M. S. Otegui, E. Panteris, C. Rasmussen, A.-C. Schmit, J. Šamaj, L. Samuels, L. A. Staehelin, D. Van Damme, G. Wasteneys, V. Žárský, Plant Cytokinesis: Terminology for Structures and Processes. Trends Cell Biol. 27, 885–894 (2017).

2. T. Murata, T. Sano, M. Sasabe, S. Nonaka, T. Higashiyama, S. Hasezawa, Y. Machida, M. Hasebe, Mechanism of microtubule array expansion in the cytokinetic phragmoplast. Nat. Commun. 4, 1967 (2013).

3. Y.-R. J. Lee, Y. Hiwatashi, T. Hotta, T. Xie, J. H. Doonan, B. Liu, The Mitotic Function of Augmin Is Dependent on Its Microtubule-Associated Protein Subunit EDE1 in Arabidopsis thaliana. Curr. Biol. 27, 3891–3897.e4 (2017).

4. Y.-R. J. Lee, B. Liu, Microtubule nucleation for the assembly of acentrosomal microtubule arrays in plant cells. New Phytol. 222, 1705–1718 (2019).

5. P. Livanos, S. Müller, Division Plane Establishment and Cytokinesis. Annu. Rev. Plant Biol. (2019), doi:10.1146/annurev-arplant-050718-100444.

6. C. G. Rasmussen, M. Bellinger, An overview of plant division-plane orientation. New Phytol. (2018), doi:10.1111/nph.15183.

7. S. Müller, S. Han, L. G. Smith, Two kinesins are involved in the spatial control of cytokinesis in Arabidopsis thaliana. Curr. Biol. 16, 888–894 (2006).

8. P. Martinez, A. Luo, A. Sylvester, C. G. Rasmussen, Proper division plane orientation and mitotic progression together allow normal growth of maize. Proc. Natl. Acad. Sci. U. S. A. 114, 2759–2764 (2017).

9. A. L. Cleary, L. G. Smith, The Tangled1 gene is required for spatial control of cytoskeletal arrays associated with cell division during maize leaf development. Plant Cell. 10, 1875–1888 (1998).

10. D. Stöckle, A. Herrmann, E. Lipka, T. Lauster, R. Gavidia, S. Zimmermann, S. Müller, Putative RopGAPs impact division plane selection and interact with kinesin-12 POK1. Nat Plants. 2, 16120 (2016).

11. S.-Z. Wu, M. Bezanilla, Myosin VIII associates with microtubule ends and together with actin plays a role in guiding plant cell division. Elife. 3 (2014), doi:10.7554/eLife.03498.

12. M. Chugh, M. Reißner, M. Bugiel, E. Lipka, A. Herrmann, B. Roy, S. Müller, E. Schäffer, Phragmoplast Orienting Kinesin 2 Is a Weak Motor Switching between Processive and Diffusive Modes. Biophys. J. 115, 375–385 (2018).

13. P. Martinez, R. Dixit, R. S. Balkunde, A. Zhang, S. E. O’Leary, K. A. Brakke, C. G. Rasmussen, TANGLED1 mediates microtubule interactions that may promote division plane positioning in maize. J. Cell Biol. 219 (2020), doi:10.1083/jcb.201907184.

14. Z. Kong, T. Hotta, Y. R. Lee, T. Horio, B. Liu, The {gamma} -tubulin complex protein GCP4 is required for organizing functional microtubule arrays in Arabidopsis thaliana. Plant Cell. 22 I : 1, 191–204 (2010).

15. B. Liu, H. C. Joshi, B. A. Palevitz, Experimental manipulation of gamma-tubulin distribution in Arabidopsis using anti-microtubule drugs. Cell Motil. Cytoskeleton. 31, 113–129 (1995).

16. S. M. Wick, Immunofluorescence microscopy of tubulin and microtubule arrays in plant cells. III. Transition between mitotic/cytokinetic and interphase microtubule arrays. Cell Biol. Int. Rep. 9, 357–371 (1985).

17. E. Panteris, P. Apostolakos, B. Galatis, Telophase-interphase transition in taxol-treated Triticum root cells: cortical microtubules appear without the prior presence of a radial perinuclear array. Protoplasma. 188, 78–84 (1995).

18. D. J. Flanders, D. J. Rawlins, P. J. Shaw, C. W. Lloyd, Re-establishment of the interphase microtubule array in vacuolated plant cells, studied by confocal microscopy and 3-D imaging. Development (1990) (available at http://dev.biologists.org/content/110/3/897.short).

19. A. Boudaoud, A. Burian, D. Borowska-Wykręt, M. Uyttewaal, R. Wrzalik, D. Kwiatkowska, O. Hamant, FibrilTool, an ImageJ plug-in to quantify fibrillar structures in raw microscopy images. Nat. Protoc. 9, 457–463 (2014).

20. T. Vavrdová, O. Šamajová, P. Křenek, M. Ovečka, P. Floková, R. Šnaurová, J. Šamaj, G. Komis, Multicolour three dimensional structured illumination microscopy of immunolabeled plant microtubules and associated proteins. Plant Methods. 15, 22 (2019).

21. K. L. Walker, S. Müller, D. Moss, D. W. Ehrhardt, L. G. Smith, Arabidopsis TANGLED Identifies the Division Plane throughout Mitosis and Cytokinesis. Curr. Biol. 17, 1827–1836 (2007).

22. L. G. Smith, S. M. Gerttula, S. Han, J. Levy, Tangled1: a microtubule binding protein required for the spatial control of cytokinesis in maize. J. Cell Biol. 152, 231–236 (2001).

23. C. G. Rasmussen, B. Sun, L. G. Smith, Tangled localization at the cortical division site of plant cells occurs by several mechanisms. J. Cell Sci. 124, 270–279 (2011).

24. S. Schmidt, A. Smertenko, Identification and characterization of the land-plant-specific microtubule nucleation factor MACET4. J. Cell Sci. 132 (2019), doi:10.1242/jcs.232819.

25. T. Sasaki, M. Tsutsumi, K. Otomo, T. Murata, N. Yagi, M. Nakamura, T. Nemoto, M. Hasebe, Y. Oda, A Novel Katanin-Tethering Machinery Accelerates Cytokinesis. Curr. Biol. 29, 4060–4070.e3 (2019).

26. K. J. Lough, K. M. Byrd, C. P. Descovich, D. C. Spitzer, A. J. Bergman, G. M. Beaudoin, L. F. Reichardt, S. E. Williams, Telophase correction refines division orientation in stratified epithelia. Elife. 8 (2019), doi:10.7554/eLife.49249.

27. A. Mohanty, A. Luo, S. DeBlasio, X. Ling, Y. Yang, D. E. Tuthill, K. E. Williams, D. Hill, T. Zadrozny, A. Chan, A. W. Sylvester, D. Jackson, Advancing cell biology and functional genomics in maize using fluorescent protein-tagged lines. Plant Physiol. 149, 601–605 (2009).

28. Q. Wu, A. Luo, T. Zadrozny, A. Sylvester, D. Jackson, Fluorescent protein marker lines in maize: generation and applications. Int. J. Dev. Biol. 57, 535–543 (2013).

29. C. G. Rasmussen, Using Live-Cell Markers in Maize to Analyze Cell Division Orientation and Timing. Methods Mol. Biol. 1370, 209–225 (2016).

30. P. Thévenaz, StackReg: an ImageJ plugin for the recursive alignment of a stack of images. Biomedical Imaging Group, Swiss Federal Institute of Technology Lausanne. 2012 (1998).

31. M. Doube, M. M. Kłosowski, I. Arganda-Carreras, F. P. Cordelières, R. P. Dougherty, J. S. Jackson, B. Schmid, J. R. Hutchinson, S. J. Shefelbine, BoneJ: Free and extensible bone image analysis in ImageJ. Bone. 47, 1076–1079 (2010).

32. R. Computing, Others, R: A language and environment for statistical computing. Vienna: R

33. H. Wickham, W. Chang, Others, ggplot2: An implementation of the Grammar of Graphics. R package version 0. 7, URL: http://CRAN.R-project.org/package=ggplot2.3 (2008).

